# Predator morphology affects prey consumption: evidence from an anuran population in subtropical wetlands

**DOI:** 10.1101/2020.06.02.130187

**Authors:** Camila Maria Mendonça da Silva, Diego Anderson Dalmolin, Laura Kauer Schuck, Camila Fernanda Moser, Alexandro Marques Tozetti

## Abstract

Morphology and diet are key factors in the ecology of organisms, determining aspects of natural history and evolution of the species. In this work, we evaluated the diet-morphology relationship in an anuran population, measuring the influence of morphological traits on the variation in the diet of individuals of *Leptodactylus latrans*. For this purpose, we collected individuals from a natural grassland habitat in southern Brazil. We analyzed the stomach content of individuals and consumed food items were classified up to the level of order. We also measured four morphological traits per individual of *L. latrans*: snout-vent length, relative limb length, distance between eyes and relative mouth width. We applied Linear Mixed Effect Models to evaluate the relationship of anuran morphological traits, number of prey taxa and volume of consumed prey. We tested the hypothesis that the configuration of predator morphological traits determines both the number of taxa and the volume of consumed prey. Our results indicate that individuals of *L. latrans* with larger body size consume a larger volume of prey and mouth width is directly and positively associated with the number of consumed taxa. In the same way that body size seems to define the capacity to ingest a large number of prey items, mouth width could be a limiting factor in prey selection. The capacity to consume a large prey volume could be an advantage in unpredictable environments, especially those with great daily thermal amplitudes such as the subtropical Brazilian grasslands.

## Introduction

Amphibians are important predators of small organisms that represent a considerable portion of the biomass present in ecosystems (Wells, 2007; Rowland *et al*., 2016). The diet of most amphibian species consists almost exclusively of arthropods; however, anurans are known for having generalist eating habits, preying on a wide variety of invertebrate orders (Duellman, 2005). Among the aspects of the natural history of amphibians, diet is recognized as one of the most important ones, reflecting their evolutionary history (Duellman & Trueb, 1994; Da Rosa *et al*., 2002).

Most studies on anuran diet are based on the quantification and description of food items and their relative importance for the species’ trophic ecology (Moser *et al*., 2017; Dias *et al*., 2018; Farina *et al*., 2018; Moser *et al*., 2019; Oliveira *et al*., 2019). However, current ecological studies reinforce the importance of using approaches that integrate the species’ ecological-evolutionary (e.g., Queiroz et al., 2015; Marques *et al*., 2019; Dalmolin *et al*., 2019), which includes the relationship within functional traits (such as morphological traits) and ecological variables. Using this approach, the processes that determine the ecological-evolutionary aspects of species become clearer (McGill *et al*., 2006; Meynard *et al*., 2011; Tonkin *et al*., 2016), and this includes their eating behavior (Tozetti & Martins, 2019). Thus, the diet stands out as an important factor associated with the ecological and morphological traits of each taxon (Sih & Christensen, 2001).

A species’ foraging strategy is considered a determining factor of the diet in anurans (Toft, 1981; Santos *et al*., 2004; Piatti & Santos, 2011). Species that are active foragers tend to consume smaller prey with social habits and slow movements (such as ants and termites). On the other hand, species that have a sit-and-wait type of forage generally consume larger and solitary prey such as beetles, orthopterans and spiders (Huey & Pianka, 1981; Toft, 1981, 1985; Strüssmann *et al*. 1984; Magnusson *et al*. 1985; Pough & Taigen, 1990). However, a substantial part of the variation in diet can emerge in response to the functional divergence between individuals and species (Simon & Toft, 1991; Piatti & Santos, 2011). In these cases, not only the behavioral aspects would be important, but also those related to the species’ physiology and morphology.

In anurans, the association between morphology and diet may be detectable when individuals of the same species are compared. The morphology of the skull structures, for example, has a great influence on the consumed prey types and can vary widely within and between populations (Emerson & Bramble, 1993; Metzger & Herrel, 2005; Cvijanović *et al*., 2014). Generally, individuals with narrower jaws tend to consume smaller prey items, whereas individuals with wider jaws tend to consume larger prey (Emerson, 1985; Menzies & Parker, 2018). The predator body size is related to their ability to subdue prey and determines the maximum size of potential prey items, thus generating relevant effects on the species’ trophic niche, as well as on the patterns of divergence in the diet between individuals with different body sizes (Strüssmann *et al*. 1984; Shine, 1991; Forsman & Lindell, 1993; Araújo *et al*., 2007). Finally, this distinction between individuals of the same species when consuming prey with different sizes can be interpreted as an important solution for intraspecific competition, thus allowing co-occurrence of phylogenetically close individuals (Guimarães *et al*. 2011; Piatti & Souza, 2011).

Based on previous studies, many aspects related to the diet of predators (e.g., species composition and biomass) are affected by their morphological traits (such as body size and mouth width; Araújo et al., 2007; Solé et al., 2017; Tozetti & Martins, 2019). However, there is a lack of understanding of how these traits act in determining predator feeding ecology. In this work, we evaluated the relationship between the descriptors of feeding behavior (richness and volume of consumed prey) and the morphological traits of *Leptodactylus latrans*, a widely distributed frog in Brazil, Paraguay, Argentina and Uruguay (Maneyro *et al*., 2004). The diet of this species is considered generalist and its foraging behavior is of the sit-and-wait type. Specifically, we tested the hypothesis that predator morphological traits (snout-vent length, relative limb length, distance between eyes and relative mouth width) determine the richness and volume of prey consumption. We predicted that consumed prey richness and volume have a positive relationship with morphological traits, especially with snout-vent length and mouth width.

## Material and Methods

### Study Area

We conducted the study in natural grassland habitats located in the municipality of Tapes, state of Rio Grande do Sul, Brazil (51 ° 22’36,8’’S; 30 ° 25’58,3’’W; Fig. 1). The study site covers an area of 1294.5 hectares and includes well-preserved areas of the Pampa Biome (in particular, sandbank formations; Becker et al., 2007). The area stands out for having one of the last remnants in Brazil of palm groves formed by *Butia odorata* (Barb. Rodr.). There is an estimated number of 70,000 individuals of *Butia odorata* forming a single continuous palm grove spread over an area of 750 ha (Fig 2 of Supplementary Material). The average annual temperature and rainfall in the study region are 18°C and 1200 mm, respectively (Maluf, 2000). The area adjacent to the palm groove is used for agricultural activities and is subject to the conventional planting system. This area has a size of about 800 ha where irrigated rice and soybean are cultivated.

**Figure 1:**
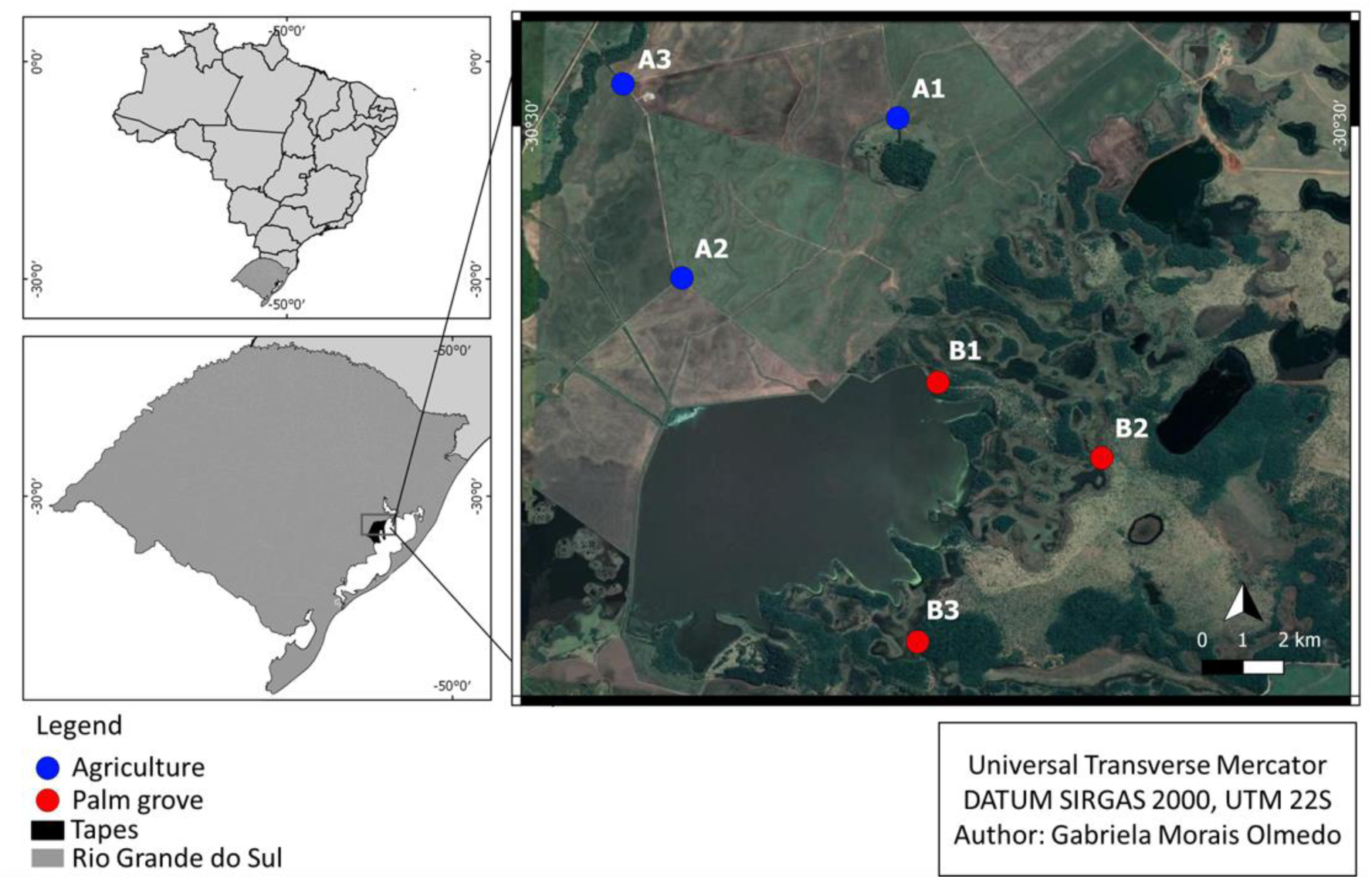
Study area in the municipality of Tapes, Rio Grande do Sul, Brazil. Sampled areas are represented by blue (agriculture) and red (palm grove) circles.

**Figure 2:**
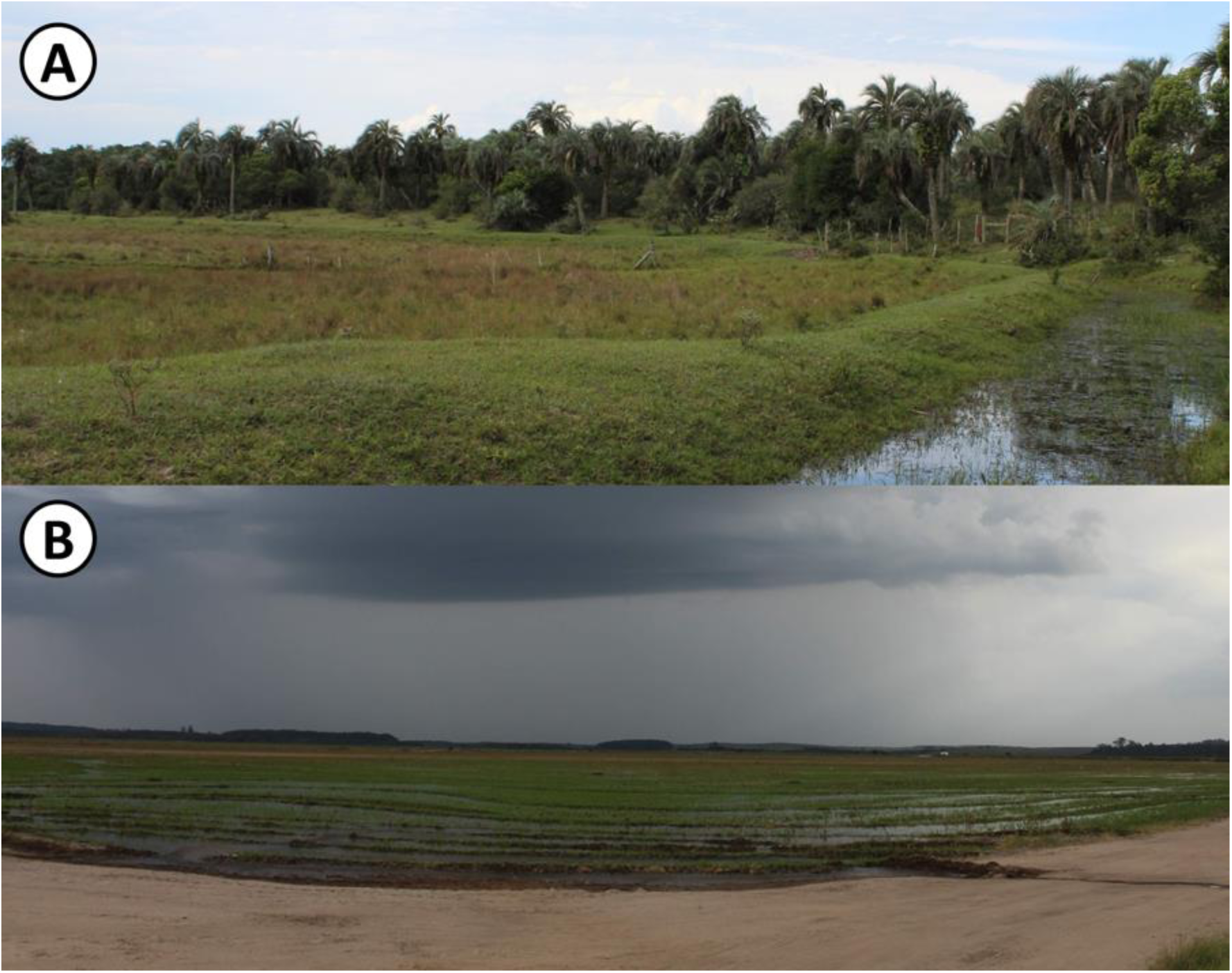
Habitat types present in the study area: (a) natural grasslands with palm groove; (b) grasslands associated with irrigated rice and soybean cultivation.

### Data Collection

We performed the sampling of individuals of *Leptodactylus latrans* from September to December 2018. For this purpose, we used visual search (Crump & Scott Jr, 1994) in breeding sites between 20 h and 02 h. We sampled each area monthly for three consecutive days (N = 12 days). The searche was performed by four researchers for one hour in as many breeding sites as possible. Thus, our total sampling effort was 288 hours of search. To avoid any effects of ontogeny in our analysis, we restricted collection to adult individuals. Search was conducted in six breeding sites distributed throughout the study area. We attempted to sample anurans from a great variety of habitats. Considering this, we selected three breeding sites in natural, preserved grasslands (in the palm groove area) and three breeding sites in grasslands adjacent to agricultural areas (irrigated rice and soybean cultivation; Fig. 2). The choice of the sampled areas occurred so that we could assess the influence of different environmental conditions on the species’ diet (Mendonça et al. unpublished data.).

The captured specimens were placed in plastic bags and kept in refrigeration equipment to decrease digestive activity until euthanasia (Moser *et al*., 2017). The collections were carried out under the authorization of the competent Federal Agency (SISBIO - authorization number 66513). Subsequently, individuals were euthanized with xylocaine, fixed with 10% formaldehyde and preserved in 70% ethanol. In the laboratory, we removed the gastrointestinal contents of each individual, which was also kept in 70% ethanol. We performed the screening of the samples using a stereomicroscope. We identified the consumed prey items up to the level of order. Afterward, we macerated and spread each individual’s stomach contents in a Petri dish. We used graph paper (1 mm x 1 mm) under the plate and measured the volume (V) and the number of orders found when categorizing the prey. The individuals of *Leptodactylus latrans* were deposited in the scientific collection of the Laboratório de Ecologia de Vertebrados Terrestres (LEVERT) of the Universidade do Vale do Rio dos Sinos (UNISINOS).

### Trait measurement

We evaluated four morphological traits of each individual: distance between eyes, mouth width, relative limb length and snout-vent length. We chose these traits based on perceived importance for determining the diet composition in several anuran species, including *Leptodactylus latrans*. The morphological traits were characterized according to the metrics presented in Fig. 3 and Table 1.

**Table 1.**
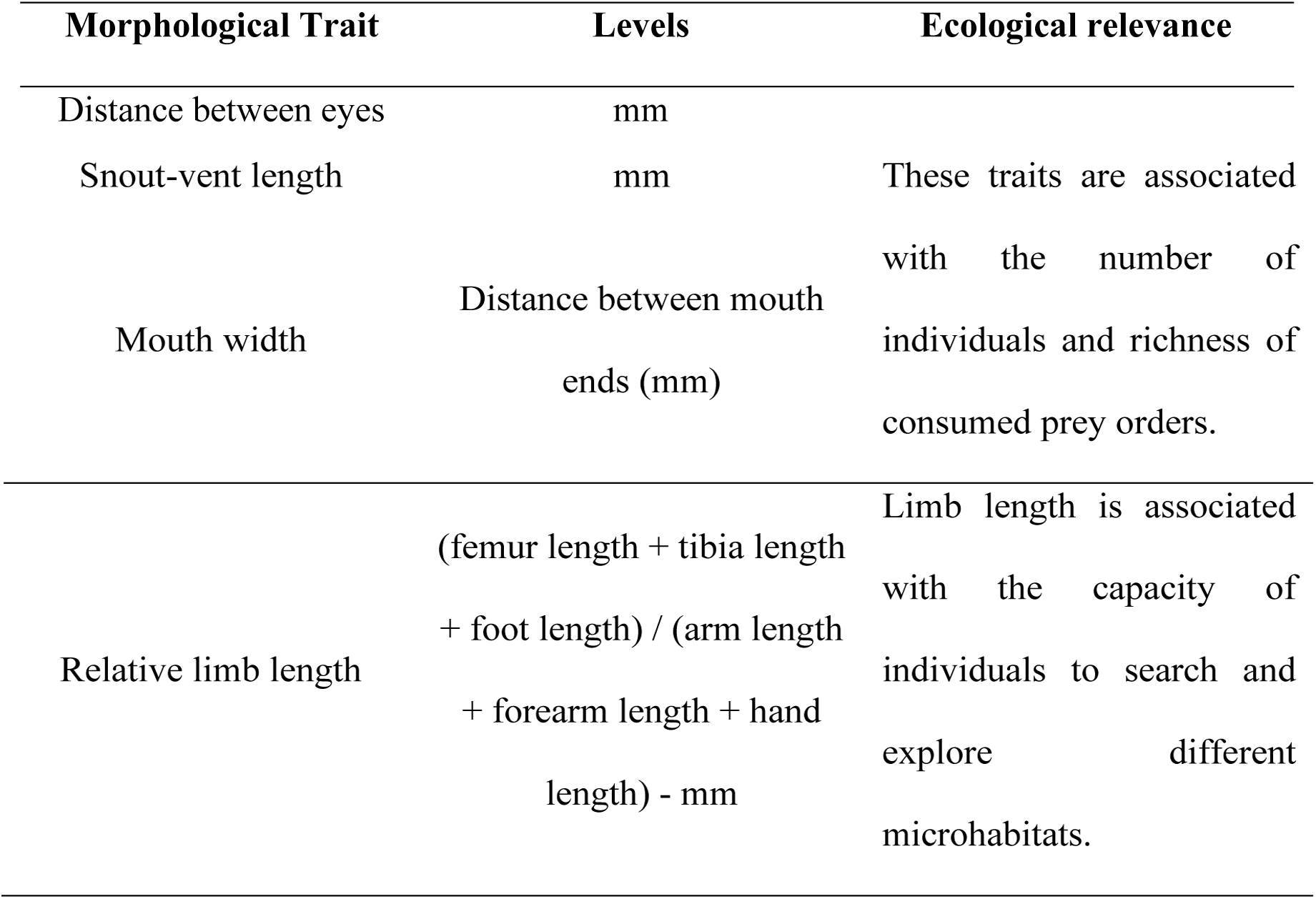
Morphological traits measured.

**Figure 3:**
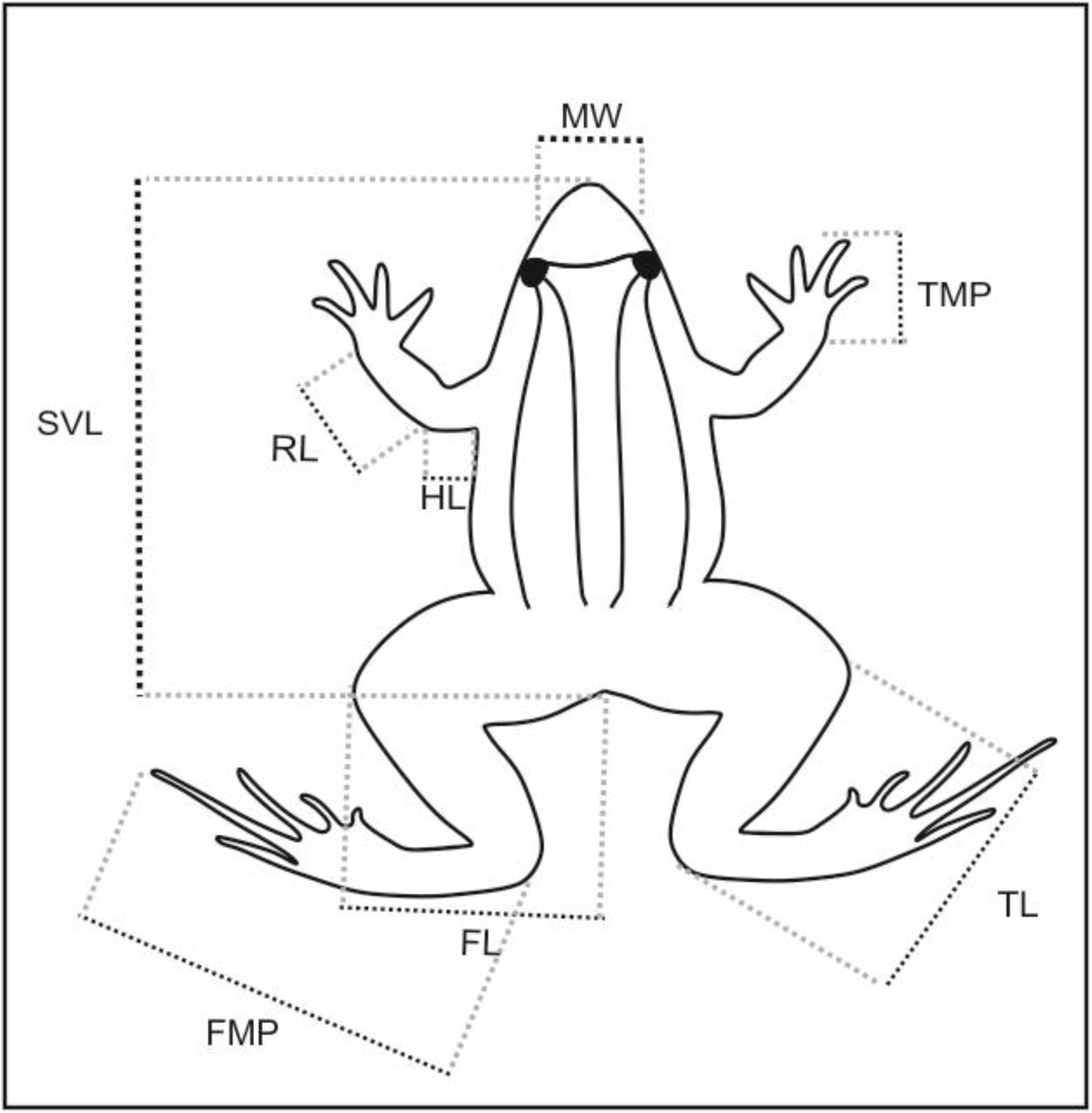
Morphological metrics evaluated in adults of *Leptodactylus latrans*: SVL = snout-vent length; TMP = third metacarpal and phalanx length; RL = radius length; HL = humerus length; FMP = fourth metatarsal and phalanx length; TL = tibia length; FL = femur length; MW= mouth width.

### Statistical Analysis

We tested the possible existence of relationships between descriptors of feeding behavior (total volume and number of consumed invertebrate orders) and morphological traits (distance between eyes, mouth width, relative limb length and snout-vent length) by LMM Poisson distribution. Date and habitat of collection were used as random effects. We log-transformed the morphological traits values before the analysis. We tested the significance of each diet-morphology relationship by ANOVA. We ran these analyses with the packages “MuMin” and “vegan” of R software.

## Results

We found a significant and positive relation of the number of consumed prey taxa and prey diversity to the mouth width (Table 2; Fig. 4a & 4b) and the total volume of consumed prey to body size (snout-vent length; Table 2; Fig. 4c). Also, the fixed effects (morphological traits) accounted for the total explained variation in both models.

**Table 2.** Results of LMM models showing the relationship between descriptors of feeding behavior and morphological traits of *Leptodactylus latrans*. R^2^m (fixed effects); R^2^C (fixed + random effects).

| Descriptors | Morphological traits | DF | *F* | *p* | AICc | R<sup>2</sup>m | R<sup>2</sup>C |
| --- | --- | --- | --- | --- | --- | --- | --- |
| **Number of invertebrate orders** | Distance between eyes | 47 | 0.25 | 0.62 |  |  |  |
|  | Mouth width | 47 | **7.53** | **0.009** |  |  |  |
|  | Relative limb length | 47 | 0.97 | 0.33 | 153.39 | 0.20 | 0.20 |
|  | Body size (snout-vent length) | 47 | 2.42 | 0.13 |  |  |  |
| **Prey diversity** | Distance between eyes | 47 | 0.99 | 0.32 |  |  |  |
|  | Mouth width | 47 | **4.02** | **0.04** |  |  |  |
|  | Relative limb length | 47 | 0.04 | 0.83 | 16.18 | 0.12 | 0.12 |
|  | Body size (snout-vent length) | 47 | 0.83 | 0.37 |  |  |  |
| **Total volume** | Distance between eyes | 47 | 0.04 | 0.85 |  |  |  |
|  | Mouth width | 47 | 1.50 | 0.23 |  |  |  |
|  | Relative limb length | 47 | 0.01 | 0.98 | 659.25 | 0.18 | 0.18 |
|  | Body size (snout-vent length) | 47 | **8.32** | **<0.001** |  |  |  |

**Figure 4:**
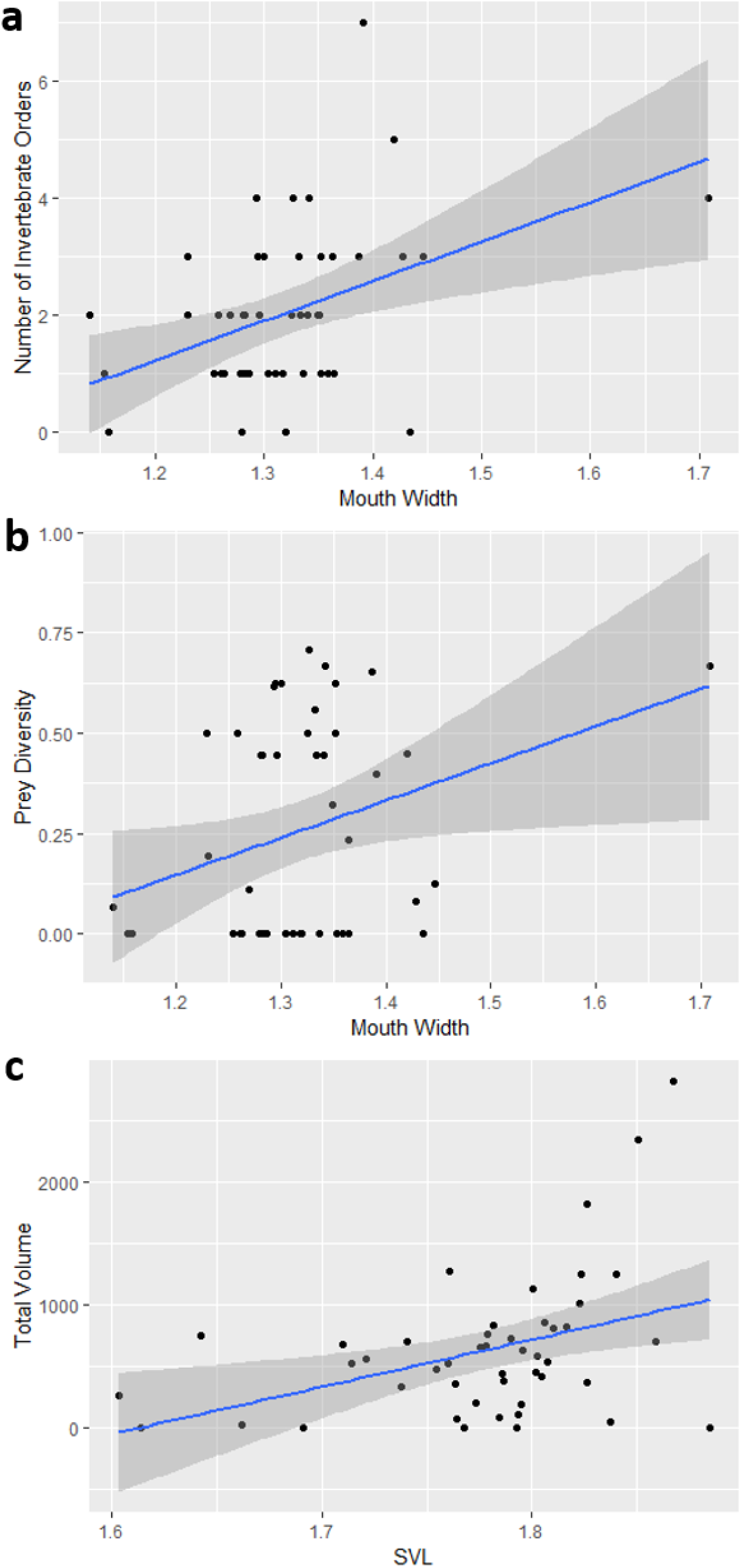
Diet-morphology relationships in *Leptodactylus latrans*. **(a)** Relationship between the number of consumed invertebrate orders and mouth width; **(b)** Relationship between the consumed prey diversity and mouth width; **(c)** Relationship between total volume of consumed prey and snout-vent length.

## Discussion

Our results indicate that the feeding habits of *Leptodactylus latrans* are influenced by an individual’s morphological traits. Our data showed that the morphological differences between individuals are reflected in variations in the volume of consumed prey and the composition of these prey items in *L. latrans*. In addition, the morphological traits affected differently each of the evaluated components (prey volume and composition). Our results suggest that: (i) individuals with larger body size (snout-vent length) consume a larger volume of prey; (ii) body size (snout-vent length) does not affect the diet composition and; (iii) the larger an individual’s mouth width, the more diverse its diet composition will be (larger number of prey taxa in the gut). These results are relevant because they highlight a little-explored dimension regarding anuran diet: intraspecific variations.

In general, species of Leoptodactylidae are considered generalist predators since they consume a great variety of food items (Piatti & Santos, 2011; Protázio *et al*., 2015). In fact, this is a common definition for many species whose diet is known (Rodrigues *et al*., 2004; Pazinato *et al*., 2011; Sugai *et al*., 2012; Dias *et al*., 2018). However, we draw attention to the fact that variations in eating habits, even subtle ones, can be found when examining the diet at the individual level. In *L. latrans*, for example, we observed that body size (snout-vent length) is one of the traits related to dietary variations. This relationship is somewhat intuitive since larger animals would supposedly have greater prey storage capacity in their digestive tracts (Tozetti & Martins, 2019). The stomach size of animals generally increases with body size, so that larger individuals tend to hold a larger volume of prey (Sugai *et al*., 2012). This hypothesis is reinforced by studies in Uruguayan populations of *L. latrans* (Maneyro *et al*., 2004), which recorded a positive correlation between the body size and the size of consumed prey. On the other hand, Solé *et al*. (2009) evaluated the diet of the same species in northeastern Brazil and found no significant correlation between the size of individuals and the volume of consumed prey. These contradictions can be attributed to methodological differences between the studies.

Our data allow us to infer that dietary patterns of *L. latrans* are determined by mouth size rather than body size. While the latter affects the “prey subduing” of consumed prey, mouth size (or its width) acts as a limiting factor in prey selection (Araújo *et al*., 2007; França *et al*. 2004; Maneyro *et al*., 2004). Limiting the opening of the mouth restricts the size and, consequently, the variety of prey that can be ingested. Still, the trend is that the relationship between mouth width and the diversity of consumed prey is linear and positive since animals with a larger mouth opening can consume a wide range of prey categories, which would contain prey items of all sizes (De Carvalho Batista *et al*., 2011; Sales *et al*., 2011). These hypotheses are reinforced by the fact that our data indicate mouth width as the main predictor of prey diversity.

Previous studies have also shown a positive relationship between morphological traits related to the head (for example, head size) and the total volume and diversity of prey consumed by anurans (Toft, 1981; Dietl *et al*., 2009). Individuals with larger heads have a greater mouth opening and this leads to an increase in the trophic niche breadth (Toft, 1981; Dietl *et al*., 2009). The fact that we did not find a relationship between limb length and diet suggests that only specific morphological traits related to foraging affect eating behavior. In fact, relative limb length does not seem to have a major contribution to foraging in anurans (Bredeweg *et al*., 2019), although this trait is a conditioning factor for access to habitats.

The relationship between a predator’s body size and the amount of prey it eats is an ecological consequence of feeding ecology (MacArthur & Pianka, 1966; Charnov, 1976; Duelmann & Trueb, 1994). The possibility of ingesting a larger volume of prey can be an important attribute in unpredictable environments and environments subjected to anthropic modification, especially those with great daily thermal amplitudes (e.g., agricultural areas). In this type of environment, sudden drops in temperature can decrease the activity of invertebrates, reducing prey availability for anurans. In this case, larger individuals would have accumulated reserves in their stomachs from nights of successful foraging.

Our data reinforce the need to consider aspects of the functional diversity of anurans (especially those related to their morphology) in future studies that are interested in the diet and trophic ecology of this group. Given the fact that anurans have high degrees of phylogenetic conservation in several traits (including morphological and behavioral traits; Campos *et al*., 2019), we suggest that these patterns of trait-diet relationship can be extended to other species of the Leptodactylidae clade.

