## Supplementary files for "Predator morphology affects prey consumption: evidence from an anuran population in subtropical wetlands"

**Supporting Information**

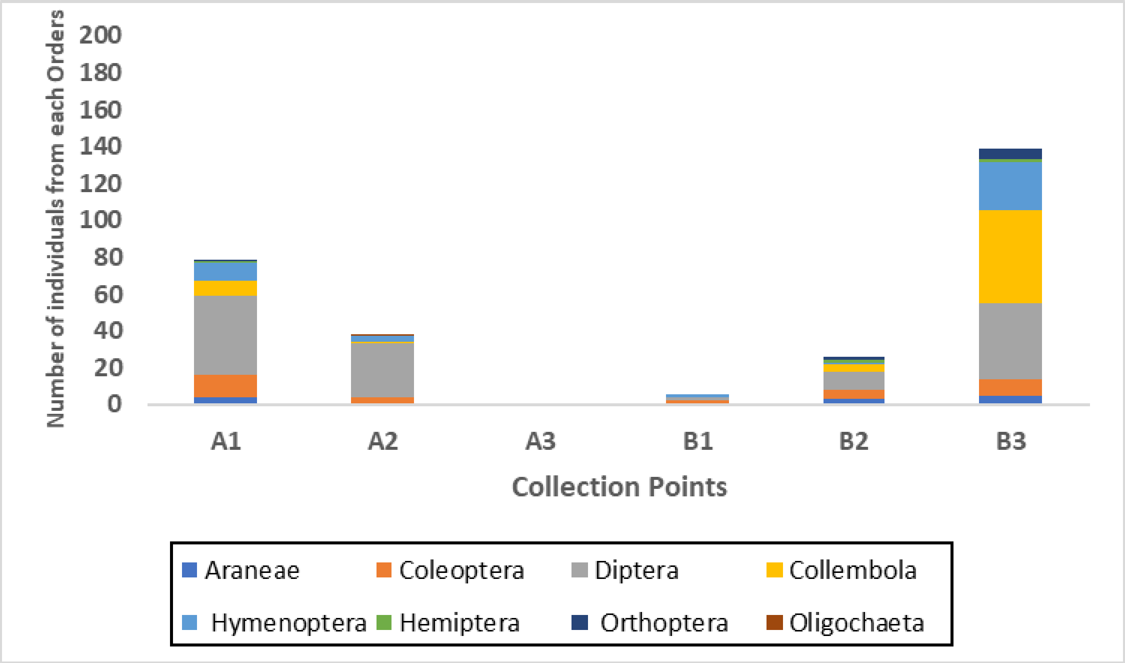


**Figure S1** – Prey availability in the six study areas assessed from September to October 2018.

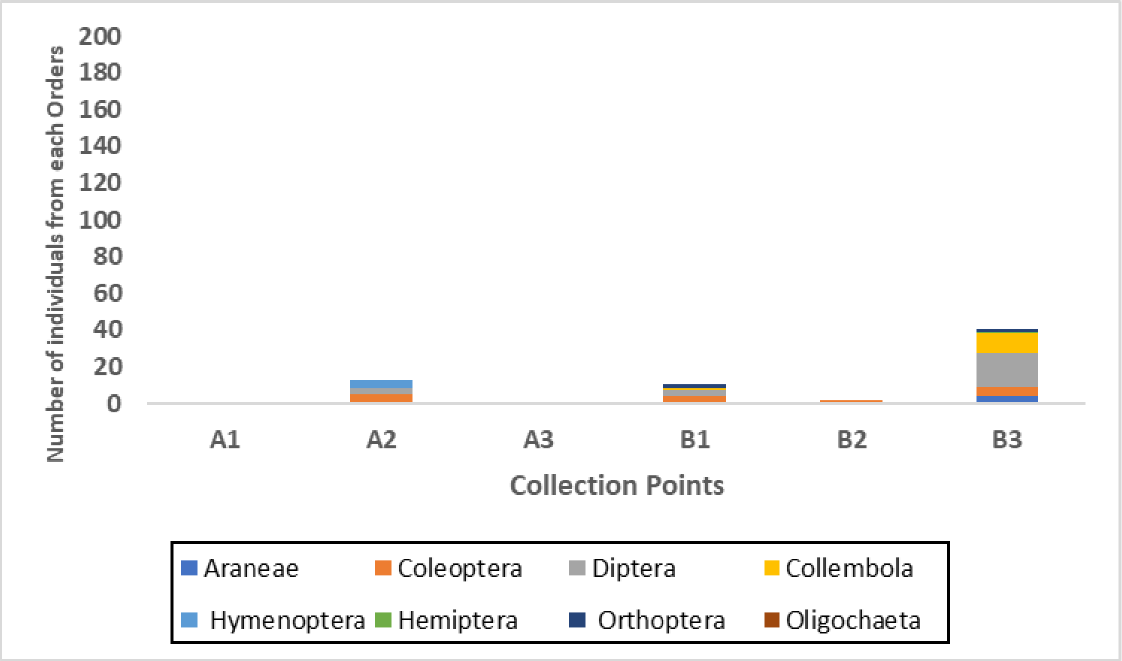


**Figure S2** – Prey availability in the six study areas assessed from October to November 2018.

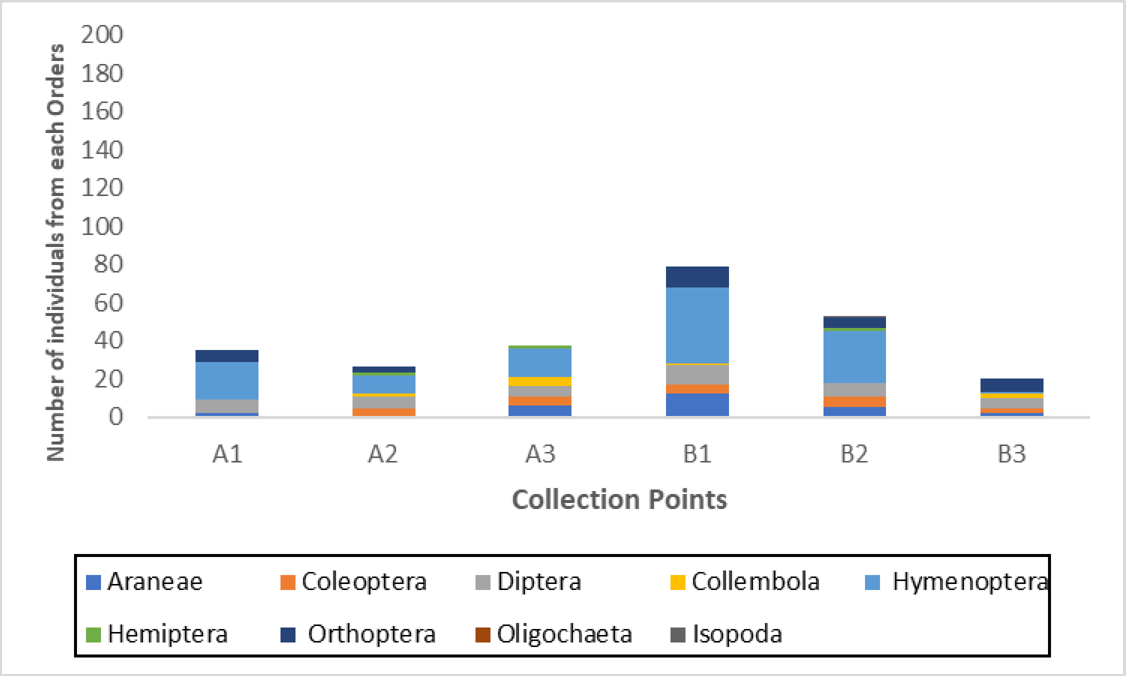


**Figure S3** – Prey availability in the six study areas assessed from November to December 2018.

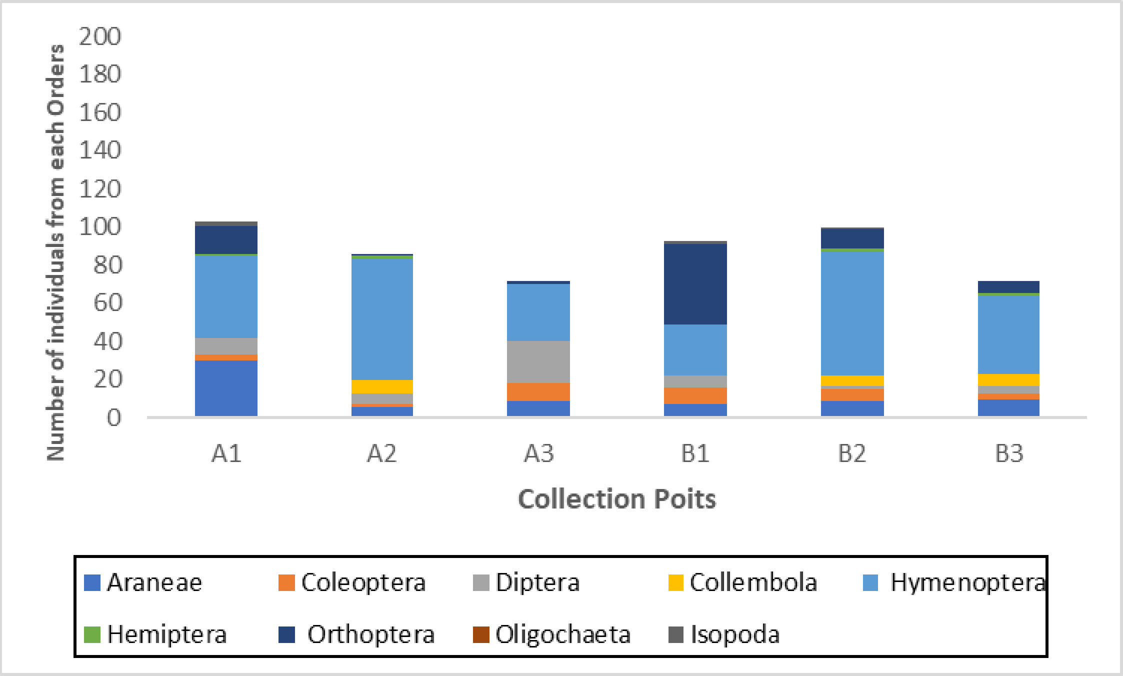


**Figure S4** – Prey availability in the six study areas assessed from December 2018 to January 2019.
